# Single-cell transcriptomic profile reveals macrophage heterogeneity in medulloblastoma and their treatment-dependent recruitment

**DOI:** 10.1101/2020.02.12.945642

**Authors:** Mai T. Dang, Michael Gonzalez, Krutika S. Gaonkar, Komal S. Rathi, Patricia Young, Sherjeel Arif, Li Zhai, Md Zahidul Alam, Samir Devalaraja, Tsun Ki To, Ian W. Folkert, Pichai Ramen, Jo Lynne Rokita, Daniel Martinez, Jaclyn N. Taroni, Joshua Shapiro, Casey S. Greene, Candace Savonen, Hakon Hakonarson, Tom Curran, Malay Haldar

## Abstract

The role of macrophages in medulloblastoma, the most common malignant pediatric brain tumor, is unclear. Using single-cell RNA sequencing in a mouse model of sonic hedgehog medulloblastoma and analysis of bulk RNA sequencing of human medulloblastoma, we investigated macrophage heterogeneity. Our findings reveal differential recruitment of macrophages with molecular-targeted versus radiation therapy and identify an immunosuppressive monocyte-derived macrophages following radiation treatment of mouse medulloblastoma, uncovering potential strategies for immunomodulation as adjunctive therapy.

## Main Text

Macrophages in the brain tumor microenvironment are emerging as a predictor of clinical outcome^1,2^. However, targeting macrophages for immunotherapy, or indeed any immunotherapy, has yet to be proven effective for brain tumor treatment. A major barrier is our incomplete understanding of the heterogeneity of tumor-associated macrophages (TAMs) and how they respond to treatment. Within tumors, anti-inflammatory (M2-polarized) macrophages drive immunosuppression while pro-inflammatory (M1-polarized) macrophages support anti-tumor immunity. Macrophages in normal tissue are phenotypically, functionally, and ontologically heterogeneous. It is currently unclear whether tumor-associated macrophages display similar heterogeneity and if insights into the nature of their heterogeneity can inform their variable functions.

Medulloblastoma (MB), one of the most common pediatric brain malignancies, is biologically heterogeneous comprising of four major molecular subgroups: WNT, sonic hedgehog (SHH), group 3, and group 4. Treatment includes surgical resection, chemotherapy, and radiation. Outcome is dependent on clinical-pathological features, patient age, and the presence of metastases. Medulloblastomas are radiosensitive but radiation can lead to severe neurological side-effects which limits its use in young children. The SHH subtype dominates this early age group and is characterized by genetic alterations leading to constitutive activation of the SHH pathway. *PTCH1* is a negative regulator of SHH signaling that is mutated in a subset of human SHH-MB patients. Germline deletion of *Ptch1* in mice leads to MB with incomplete penetrance, but this increases to 100% penetrance when combined with loss of *Tp53*^3^. Importantly, *Tp53* loss in human SHH-MB is associated with poor prognosis^4^. Hence, *Ptch1*^*+/-*^:*Tp53*^*-/-*^ mice are useful for studying high-risk SHH-MB. Inhibitors of the SHH pathway, such as GDC-0449 (Vismodegib™), have shown remarkable efficacy in treating murine SHH-MB and human patients in ongoing clinical trials^5,6^. However, development of therapeutic resistance to GDC-0449 remains a concern and GDC-0449 is contraindicated in young children because of its negative impact on bone growth^7,8^. Hence, novel therapies are vitally needed for patients in this high-risk group.

Among the four different MB subtypes, SHH-MB harbors the most macrophages^9^. The vast majority of studies on macrophages in brain tumors are based on immunohistochemistry, flow cytometry, or transcriptional profiling of bulk tumor tissue, methods that have limited capacity to resolve cellular heterogeneity^9–11^. Single-cell RNA sequencing (scRNA-Seq) overcomes this limitation and has helped uncover TAM heterogeneity in other solid tumors. In a recent study, scRNA-Seq of human MB provided new insights into tumor heterogeneity but very few macrophages were captured, precluding detailed analyses of TAMs^12^. Furthermore, it remains unclear how these macrophages respond to standard treatment. In this study, we examine TAM composition in MB by performing scRNA-Seq of TAMs in *Ptch1*^*+/-*^: *Tp53*^*-/-*^ mice and corresponding comparisons through the OpenPBTA project to estimate their contributions to bulk RNA-Seq measurements. Importantly, we identify unanticipated differences in the TAM composition in response to radiation and molecular-targeted therapy with GDC-0449, exposing potential strategies for immunomodulation as adjunctive therapy in MB.

Consistent with previous reports, we find that SHH-MB contains significantly more macrophages than surrounding normal brainstem tissue (Figure 1a). To study macrophage heterogeneity, we performed scRNA sequencing (10X genomics) on CD11b+ myeloid cells isolated from the following three types of samples from *Ptch1*^*+/-*^: *Tp53*^*-/-*^ mice: 1) cerebella of 2-week old mice with minimal tumor to capture microglia in normal tissue, 2) peripheral blood of the same mice, 3) and cerebella of 8-week old mice harboring large tumors. Analysis across all three sample types revealed significant alteration in cerebellar macrophage composition in the presence of tumor (Figure 1b,c). Two large TAM clusters (collectively denoted TAM1 henceforth) aggregated closer to microglia while another three clusters (collectively denoted TAM2 henceforth) aggregated closer to monocytes, suggesting heterogeneous cell-of-origin for macrophages in MB (Figure 1d). Monocle-based analysis also supported microglia and monocyte origins for TAM1 and TAM2 respectively (Figure 1e).

**Figure 1.**
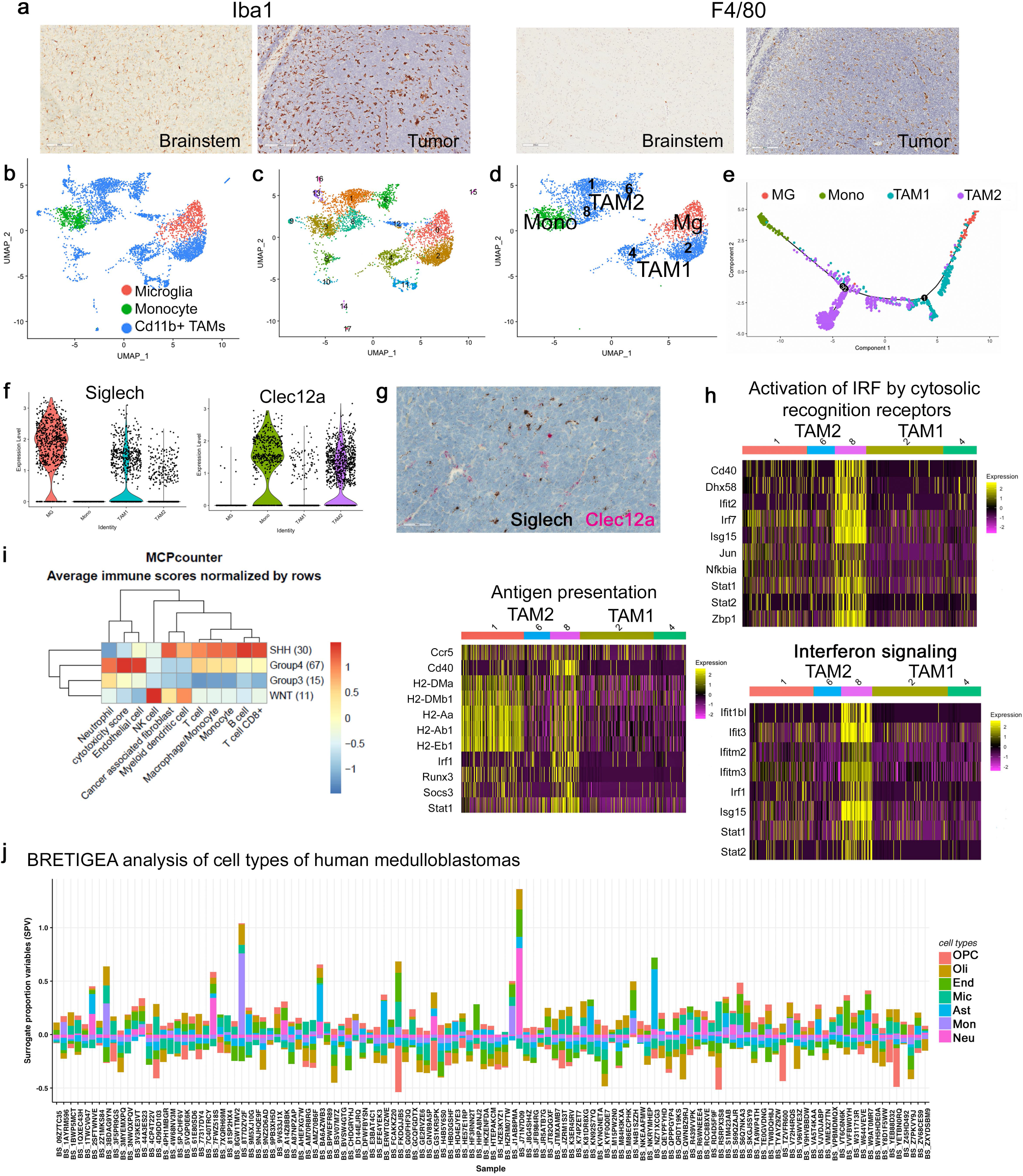
Dual origins of medulloblastoma-infiltrating macrophages. (a) Immunohistochemistry of *Ptch1*^*+/-*^:*Tp53*^*-/-*^ SHH-MB with Iba1 and F4/80 antibodies show tumor has a large accumulation of macrophages in comparison to brain stem. (b) Uniform Manifold Approximation and Projection (UMAP) display by sample type and by Seurat-based clusters (c) of cerebellar microglia and selected monocyte population from the peripheral blood of 2 week old *Ptch1*^*+/-*^:*Tp53*^*-/-*^ mice (n=3) and TAMs from cerebella of eight week old mice (n=2). (d) UMAP of select large clusters that we designate TAM1 for clusters more similar to microglia and TAM2 for those more similar to monocytes. (e) Monocle-based pseudotime analysis of the scRNA-Seq data show TAM1 being more similar to microglia and TAM2 to monocytes. (f) Violin plots displaying expression of genes known to be differentially expressed in ontologically distinct brain macrophages. Microglia-associated *Siglech* is highly expressed in TAM1 while monocyte-associated *Clec12a* is highly expressed in in TAM2. (g) RNA in situ hybridization (RNAscope™)-based detection of *Siglech* and *Clec12a* expression. The overwhelming majority of TAMs express one or the other marker, but not both. (h) IPA™ analyses of differentially expressed genes between TAM1 and TAM2. Overall, TAM2 displays higher ‘inflammatory signature’ based on enrichment of pathways associated with: antigen presentation, interferon signaling, and activation of interferon-regulated factors induced by cytosolic pattern-recognition receptors. (i) MCP-counter-based analysis of cellular composition using bulk RNA-Seq of 123 human pediatric medulloblastomas. SHH-MB show higher infiltration with lymphocytes and myeloid cells compared to other MB subtypes. (j) BRETIGEA cell proportion analysis of aforementioned human medulloblastoma RNA-Seq data show the co-existence of both microglia (Mic) and monocytes (Mono) in human MB. OPC = oligodendrocyte precursor cells, Oli = oligodendrocytes, Mic = microglia, End = endothelial cells, Ast = astrocytes, Mono = monocytes, Neu = neurons.

Previous studies have shown that only a few markers distinguish microglia from monocyte-derived macrophages in the brain and non-medulloblastoma high grade tumors^1,10,13–15^. Consistent with this, we find only a handful of markers (for example, *Siglech, Phgdh*, and *Pmp22*) that distinguished microglia in MB (Supplemental Figure 1a). Notably, we found expression of certain microglia-associated genes (for example, *Cd81, Fcrls, Olfml3, Sparc*, and *Tmem119*) in monocyte-derived TAM2 (Supplementary Figure 1b), which likely reflects the impact of the brain microenvironment on monocyte-derived macrophages. Likewise, cross-expression of certain markers (for example, *Axl, Cst7, Cxcl16*, and *Ms4a7*) in both TAM1 and TAM2 may reflect the impact of tumor-microenvironment on macrophages (Supplementary Figure 1c). Certain canonical markers for monocytes such as *Ly6c2* and *Ccr2* were preserved in TAM2 and not present in TAM1 as expected (Supplement Figure1d). Additional monocyte-associated markers that were preserved in TAM2 includes *Plac8, Tgfbi, Iqgap1, Crip1, Vlm, Sirpb1c, S100a6, Plbd1*, and *Pla2g7*. Other markers such as *Mgst1* and *Chil3* were expressed in monocytes but lost in TAM2 (Supplement Figure1e). These findings will inform ongoing and future efforts to distinguish TAMs and resident brain macrophage subsets.

We selected *Siglech* and *Clec12a* as specific markers of microglia- and monocyte-origins respectively for further validation in MB (Figure 1f). Given the vagaries of antibody-based detection, we used RNAscope™ *in situ* analysis, which confirmed expression of *Siglech* and *Clec12a* in the tumor microenvironment (Supplement Figure 2a,b). Both markers were only expressed in Iba1-positive cells in the tumor, confirming their specific expression in macrophages (Supplement Figure 2c,d). Dual staining showed mutually exclusive pattern of expression of these markers in tumor (Figure 1g). Examining TAM1 and TAM2 using ingenuity pathway analysis (IPA) showed enrichment of pathways associated with antigen presentation, interferon signaling, and activation of interferon regulated factors (IRF) downstream of cytosolic pattern-recognition receptors in TAM2 (Figure 1h).

To examine the relevance of our findings in murine SHH-MB to human patients, we used deconvoluted bulk RNAseq data from 123 patients generated by the collaborative OpenPBTA project. MCP-counter analyses^16^ of this dataset showed SHH subtype MB to be significantly enriched (amongst other MB subsets) for monocytes and macrophages, which is consistent with prior published IHC studies^10^ (Figure 1i). Additional analysis using BRETIGEA^17^ suggest that all subtypes of MB contain both microglia and monocyte-derived TAMs, albeit at varying proportions (Figure 1j, Supplementary Figure 3).

There is a great deal of interest in the oncology community to characterize immune responses to traditional forms of treatment such as radiation, chemo, or molecular-targeted therapy. The overarching goal is to identify opportunities to combine these existing treatment modalities with emerging immunotherapeutic approaches. Therefore, we next investigated TAM responses to two distinct treatment modalities; SHH-pathway inhibitor (GDC-0449, twice daily doses of 100mg/kg for 4 days) and radiation (3×10Gy). Radiation inducing a much greater macrophage recruitment (Figure 2a). We performed scRNA-Seq on CD11b+ myeloid cells isolated from tumor-harboring cerebellum of mice treated with each treatment modalities (Figure 2b,c). Using TAM1- and TAM2-associated markers identified above, we found significantly higher recruitment of monocyte-derived TAMs with radiation when compared to GDC-0449 (Figure 2d). We further validated this finding within tumors via RNAscope™ with *Siglech* and *Clec12a* probes (Figure 2e). Hence, molecular-targeted therapy with GDC-0449 and radiation therapy leads to recruitment of ontologically distinct subsets of macrophages.

**Figure 2.**
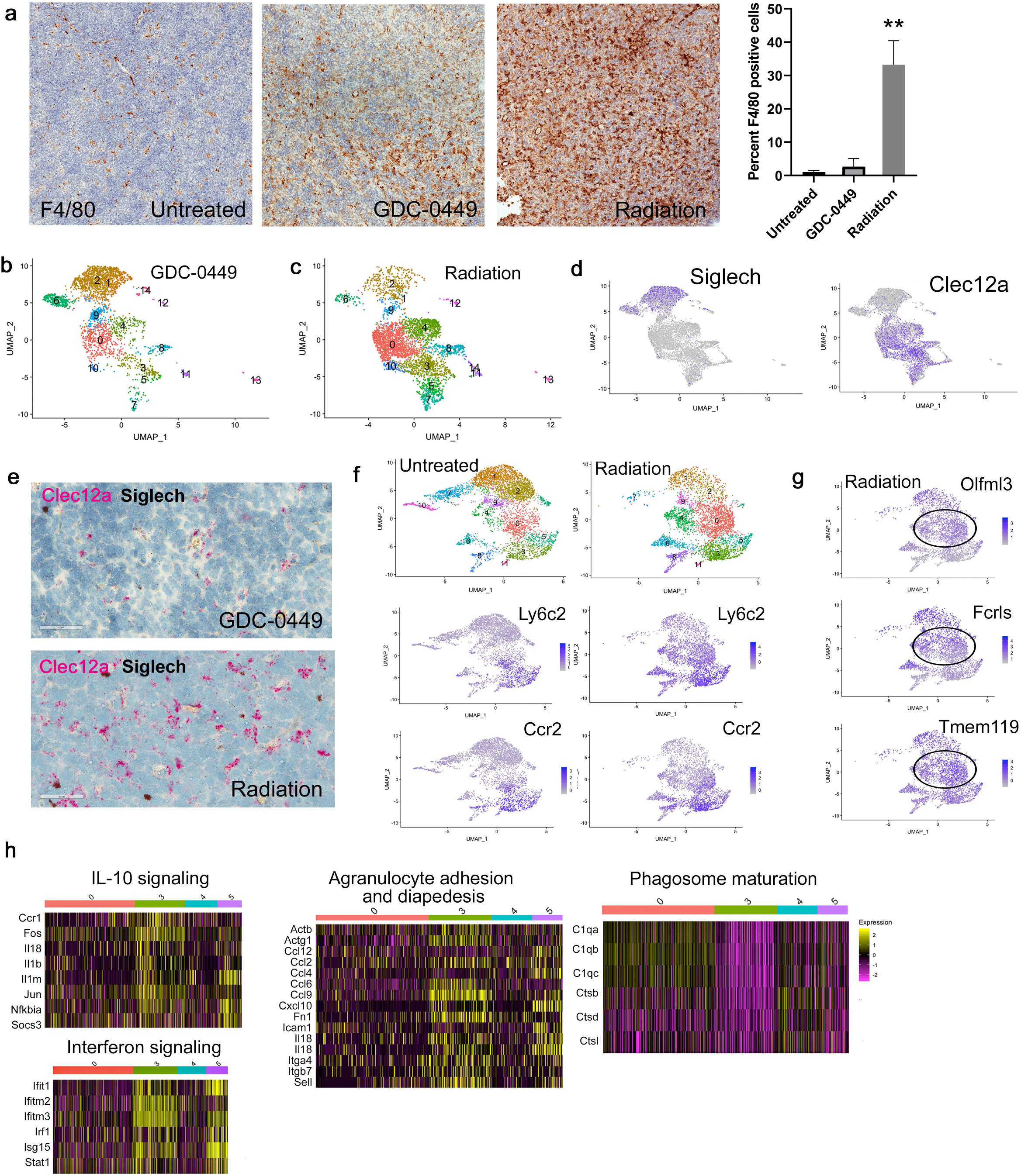
Radiation induces selective recruitment of monocyte-derived macrophages compared to molecular-targeted treatment in SHH-MB. (a) Both GDC-0449 and radiation recruits macrophages within SHH-MB with the latter displaying significantly greater recruitment. (b-c) UMAP display of aggregated data of GDC-0449 treated (n=2) and radiation treated tumors (n=2) with Seurat-based clustering show differential recruitment of TAM populations as determined by their expression of aforementioned markers for microglia and monocyte-derived TAMs, including *Siglech* and *Clec12a* (d). (e) RNA-Scope™ analyses of tumor tissue is consistent with scRNA-Seq, showing higher accumulation of *Clec12a*-expressing TAMs with radiation therapy. (f) UMAP display of scRNA-Seq comparing untreated tumors to radiation treated tumors with Seurat-based clustering (labeled by numbers). Radiation induces accumulation of TAMs expressing monocyte-markers Ly6c2 and Ccr2. (g) Among the four largest monocyte-derived clusters that showed highest increase post-radiation, clusters 0 and 4 have less Ly6c2 and Ccr2 (*Lyc2/Ccr2*^lo^, genes associated with monocytes) expression and higher expression of microglia-associated markers such as *Olfml3, Fcrls, and Tmem119*. (h) IPA of these four clusters in irradiated samples shows *Lyc2/Ccr2*^hi^ clusters (clusters 3 and 5) have higher expression of interferon signaling genes and granulocyte adhesion/ diapedesis genes, while *Lyc2/Ccr2*^lo^ clusters have higher expression of markers for phagocytosis. ** *p* = <0.01, *** *p =* <0.0001

Given the robust recruitment of TAM2 with treatment, we further analyzed this population with and without radiation. Clusters 3 and 5 within TAM2 showed higher expression of prototypical monocyte-associated genes *Ly6c2* and *Ccr2* (*Ly6c2/Ccr2*^hi^) when compared to clusters 0 and 4 (*Ly6c2/Ccr2*^*lo*^, Figure 2f). These latter clusters had higher expression of genes typically associated with microglia (*Tmem119, Fcrl2*, and *Olfml3*, Figure 2g). IPA-based comparison of these all four of these clusters showed adhesion and diapedesis activity as well as higher interferon signature in *Ly6c2/Ccr2*^hi^ subset (Figure 2i) in comparison to the *Ly6c2/Ccr2*^*lo*^. Importantly, they also show enrichment of genes in the IL-10 pathway suggesting an immune suppressive phenotype. In contrast, *Ly6c2/Ccr2*^*lo*^ show increased expression of complement and cathepsin genes, suggestive of higher phagocytic activity (Figure 2i). This may reflect a maturation spectrum where newly generated monocyte-derived macrophages may be more immune-suppressive and gradually turn on microglia associated genes in response to the brain microenvironment as they take on phagocytic functions that are characteristic of mature macrophages.

To examine the function of monocyte-derived TAM2 in MB, we generated a monocyte-deficient SHH-MB model by breeding *Ptch1*^*+/-*^: *Tp53*^*-/-*^ mice with *Ccr2* knockout (deficient in circulating monocytes) mice (Figure 3a). Henceforth, we refer to *Ptch1*^*+/-*^: *Tp53*^*-/-*^ *Ccr2*^*-/-*^ mice as Ccr2KO and *Ptch1*^*+/-*^: *Tp53*^*-/-*^ mice as Ccr2WT. We performed scRNA-Seq on immune cells isolated from tumor-bearing cerebellum from Ccr2KO and Ccr2WT mice (Figure 3b-d). Monocyte deficiency was associated with altered myeloid cell recruitment post-radiation, including a significant reduction in *Clec12* positive TAMs (Supplement Figure 4), with a pronounced reduction in *Ly6c2/Ccr2*^hi^ clusters (Figure 3e) in the Ccr2KO. This was also confirmed with flow cytometry (Figure 3f). Intriguingly, monocyte deficiency increased myeloid clusters 1 and 5, which expressed high levels of neutrophil markers such as *S100a9* and *Retlng* (Figure 3g). Consistent with this observation, we found significant accumulation of neutrophils (via Ly6G immunohistochemistry) within post-irradiated tumors of Ccr2KO compared to Ccr2WT tumors (Figure 3h).

**Figure 3.**
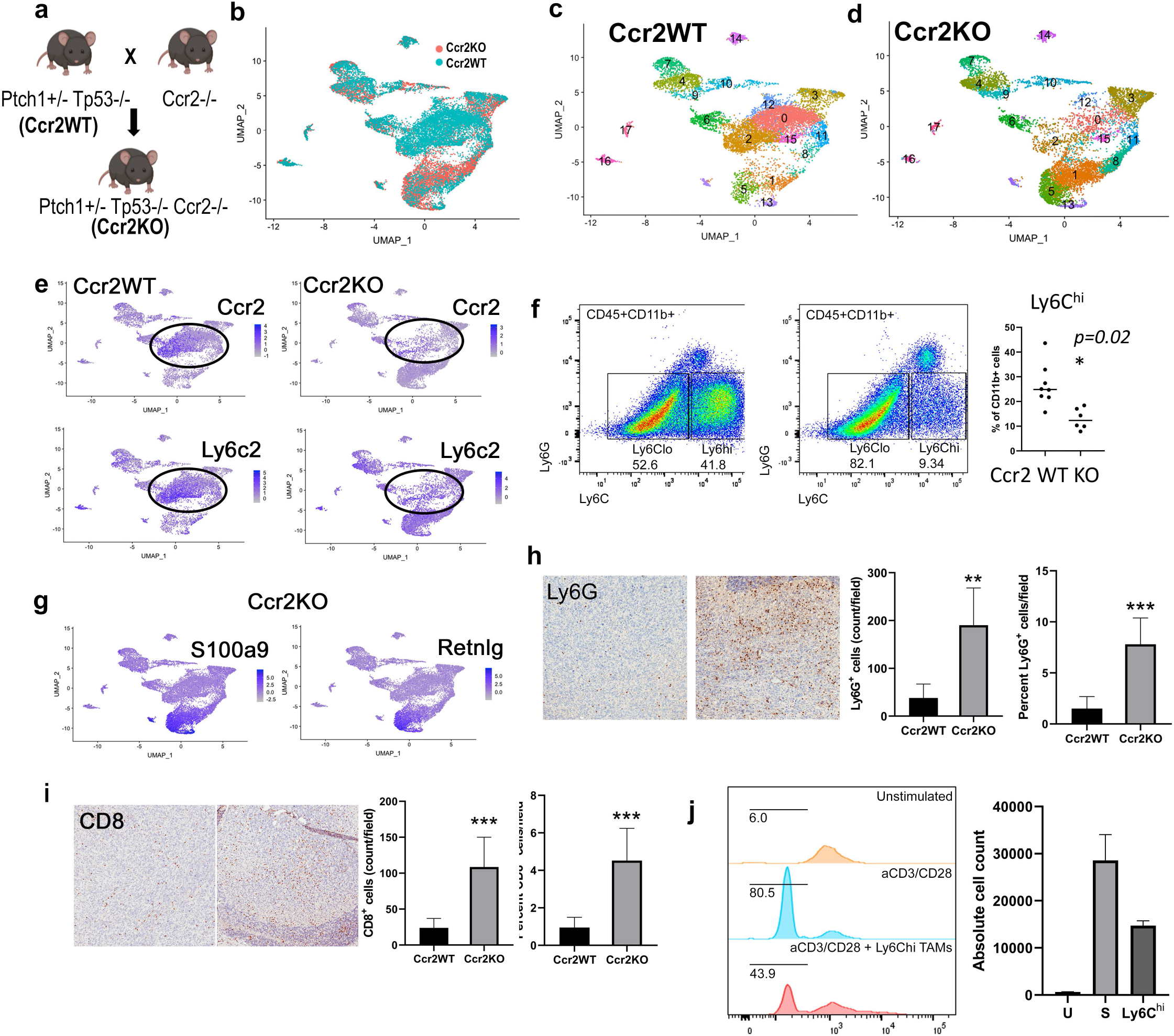
Reducing monocyte-derived macrophages engenders inflammatory signature in radiation-treated SHH-MB. (a) *Ptch1*^*+/-*^:*Tp53*^*-/-*^ (Ccr2WT) and *Ccr2*^*-/-*^ mice were bred to produce *Ptch1*^*+/-*^:*Tp53*^*-/-*^:*Ccr2*^*-/-*^ mice (Ccr2KO). (b) Aggregated analysis scRNA-Seq from irradiated SHH-MB harboring cerebellum from Ccr2WT (n=3, blue) and Ccr2KO (n=3, red) mice, grouped by genotype of Ccr2WT and Ccr2KO. Two Ccr2KO samples and one Ccr2WT sample contained cells enriched by CD45+ selection and the remainder were of Cd11b+ enriched cells. Subsets of just Ccr2WT (c) and Ccr2KO (d) cells show differential recruitment of distinct myeloid populations. Seurat-based clusters are labeled as numbers. (e) *Ly6c2/Ccr2*^hi^ subpopulations (highlighted) are significantly reduced in post-radiation Ccr2KO tumors compared to Ccr2WT. (f) Flow cytometry confirms the reduction of *Ly6c*^hi^ cells in Ccr2KO. (g) Largest myeloid subpopulation selectively recruited in Ccr2KO tumors have signature of neutrophils (*S100a9, Retnlg*). (h) Immunohistochemistry staining with Ly6G antibody shows irradiated Ccr2KO have a significantly higher accumulation of neutrophils. (i) Staining with CD8+ antibody shows they also have an increased accumulation of cytotoxic CD8+ T cells.(j) T cell proliferation assay shows *Ly6c*^hi^ TAMs from irradiated Ccr2WT tumors suppress proliferation of CD3/CD28 stimulated splenic CD8+ cells. ** *p < 0.01*

The function of neutrophils in tumor is unclear and may themselves be heterogeneous, but neutrophil infiltration is commonly associated with inflammation, suggesting that the absence of TAM2 may lead to a more inflammatory milieu in radiation treated SHH-MB. Hence, we next asked whether loss of TAM2 with monocyte deficiency might also enhance frequency of intratumoral CD8 T cells. IHC with CD8+ antibody showed significantly higher levels of CD8T cells in post-irradiated tumors in Ccr2KO tumors compared to Ccr2WT (Figure 3i). These findings suggest that monocyte-derived TAM2 are immunosuppressive. To further assess this, we performed an *in vitro* T cell proliferation assay in which we co-cultured Ly6C^hi^ cells from irradiated Ccr2WT animals with splenic T cells. Proliferation of CD8T cells were significantly inhibited in the presence of Ly6c^hi^ cells, supporting their immunosuppressive nature (Figure 3j).

Our work uncovers phenotypic and ontological heterogeneity within SHH-MB infiltrating TAMs and demonstrates distinct patterns of TAM recruitment with radiation versus molecular-targeted therapy. We find that radiation-induced monocyte derived TAMs are immunosuppressive and their absence engenders a pro-inflammatory tumor microenvironment marked by increased neutrophils and CD8+ T cells. T cell suppression by tumor-infiltrating macrophages is well-known, but the TAM interactions with neutrophils are less well understood. When compared to circulating neutrophils, we found that these tumor-associated neutrophils expressed higher levels of genes associated with communication between innate and adaptive immune system (Supplementary Figure 5). In this context, it is not entirely clear whether the increased influx of neutrophils in monocyte-deficient post-irradiated tumors are supportive or inhibitory to the tumor and/or anti-tumor immune responses and warrants further investigation. The increased frequency of CD8+ T cells suggests a potential benefit of combining immune checkpoint blockade (anti-PD1/PDL1) with monocyte depletion (Ccr2-targeting small molecule inhibitors or antibodies) in the setting of radiation therapy in MB. Given the desperate need for new treatment approaches, such evidence-based rational combination therapy may hold promise for brain tumors.

---

**Supplementary Figure 1. scRNA-Seq identifies markers for TAM subtypes and origin.** (a) Microglia-specific genes that are higher in TAM1 compared to TAM2. (b) Microglia-specific genes that are expressed in both TAM1 and TAM2, highlighting the influence of the brain microenvironment. (c) Genes selectively expressed in TAMs but not in the cell of origin, highlighting the influence of tumor microenvironment. d) Monocyte-specific genes expressed in TAM2 but not TAM1. (e) Monocyte-associated genes that are not expressed inTAM2.

**Supplementary Figure 2. Specificity of *Siglech* and *Clec12a* in tissue.** (a) *Siglech* is expressed in microglia of the brainstem (arrow points to probe positive cells) while *Clec12a* (b) does not (c). (d) *Siglech* and *Clec12a* are only expressed by IbaI^+^ cells.

**Supplementary Figure 3. Cell proportion of microglia and monocyte in tumor tissue in the 4 subtypes of human medulloblastoma.** BRETIGEA analysis of deconvoluted bulk RNAseq human medulloblastomas show varying relative proportions of microglia versus monocyte-like macrophages within subtypes. Macrophages with microglia signature are more abundant in SHH (Wilcoxon *p =* 0.026, n=48) and WNT (Wilcoxon *p* = 0.022, n=20) subtypes, while macrophages with monocyte markers are higher in Group 4 (Wilcoxon *p* = 1.6e-5, n=116).

**Supplementary Figure 4. Expression of Clec12a expression in irradiated tissue.** a) scRNA-Seq data of Ccr2WT and Ccr2KO samples post-radiation show *Clec12a* expressing TAMs are significantly reduced in post-irradiated Ccr2KO tumors compared to Ccr2WT. b) RNAscope™ using probes for Siglech (brown) versus Clec12a (red) show relatively less Clec12a expressing cells than in Ccr2WT shown in Figure 2f.

**Supplementary Figure 5. IPA identifies increase expression of genes associated with innate and adaptive immune interaction.** (a) UMAP showing S100a9 expressing neutrophils from peripheral Cd11b+ sample and (b) from irradiated Ccr2KO mice. (c) IPA shows increased expression of cytokines and MHC-associate genes, suggesting potential interactions with adaptive immune cells in post-irradiated MB-associated neutrophils compared to circulating neutrophils.

## Material and methods

### Animals

Three genetically engineered models purchased from Jackson Laboratory were used: Ptch^+/-^ (stock no. 003081), Tp53^-/-^ (stock no. 002101), Ccr2^-/-^ (004999). Ptch1^*+/*-^:Tp53^-/-^ mice were bred and Ccr2^-/-^ genotyping was performed as previously reported^3,18^. Both males and females of both Ptch1^*+/-*^:Tp53^-/-^ and Ptch1^*+/*-^:Tp53^-/-^Ccr2^-/-^ were used for this study. Mice were bred and maintained in specific pathogen free facilities at the University of Pennsylvania. Mice were group-housed (21°C; 12h:12h light:dark cycle) and given *ad libitum* access to standard rodent diet. All animal procedures were conducted according to National Institutes of Health guidelines and approved by the Institutional Animal Care and Use Committee at the University of Pennsylvania.

### Tumor treatment modalities

Mice were treated with SMO inhibitor, GDC-0449 (Chemitek, suspended in 0.5% methylcellulose and 0.2% Tween 80) at a dose of 100mg/kg twice a day for 4 days. Brains were resected and analyzed 12 hours after the 8^th^ dose. Radiation treatment was completed on a Small Animal Radiation Research Platform that delivers photon radiation with CT-guided location. Mice that were used for scRNA-Seq received 3 doses of 10Gy radiation on consecutive days. Whole brain were resected for TAM isolation 5 days after the last dose of radiation. Tissues from irradiated mice for all other experiments received 1 dose of 10Gy radiation and tissues were analyzed 3 or 4 days after radiation.

For radiation treatment, mice were anesthesized with 2% isoflurane in a carrier gas of medical grade air utilizing an induction chamber connected to anesthesia machine (Matrx). Once the mouse reached the desired plane of anesthesia (∼2 minutes), the animal was placed on the SARRP’s (Xstrahl Life Sciences) irradiation platform with the nose of the mouse in a nosecone where flow of administered isoflurane (maintained at 2%) was remotely controlled using Somnosuite (Kent Scientific) anesthesia system. The mouse tumor was targeted manually with the help of onboard positioning lasers of the SARRP. 10 Gy doses were delivered using 10 mm diameter collimated beam of X-rays with tube potential of 220kVp, 13mA current and dose rate of ∼2Gy/min.

### scRNA-Sequencing and analysis

Whole cerebellar tissue were mechanically homogenized in RPMI medium. Immune cells were purified using a 70/30 Percoll (Sigma Aldrich, 17-0891-02) gradient spun for 30 minutes at 500G. Cells at the interface were collected and further purified with CD11b+ magnetic microbeads (Miltenyi Biotec, 130-092-636) or CD45 beads (Miltenyi Biotec, 130-052-301) through two consecutive LS columns (Miltenyi Biotec, 130-042-401). Peripheral blood was collected in heparinized tubes and spun down to collect cellular components. Red blood cells were lysed (ACK lysing buffer). Remaining lymphocytes were isolated with CD11b+ magnetic beads.

Next-generation sequencing libraries and sequencing were conducted at the Center for Applied Genomics Core at the Children’s Hospital of Philadelphia. Libraries were prepared using the 10x Genomics Chromium Single Cell 3’ Reagent kit v2 per manufacturer’s instructions. Libraries were uniquely indexed using the Chromium i7 Sample Index Kit, pooled, and sequenced on an Illumina HiSeq sequencer in a paired-end, single indexing run. Sequencing for each library targeted 20,000 mean reads per cell. Data was processed using the Cellranger pipeline (10x genomics, v.3.0.2) for demultiplexing and alignment of sequencing reads to the mm10 transcriptome and creation of feature-barcode matrices. Individual single cell RNAseq libraries were aggregated using the cellranger aggr pipeline. Libraries were normalized for sequencing depth across all libraries during aggregation. Secondary analysis on the aggregated feature barcode matrices was performed using the Seurat package (v.3.0) within the R computing environment. Briefly, cells expressing less than 200 or more than 4000 genes were excluded from further analysis. Additionally, cells expressing greater than 10% mitochondrial genes were excluded from the dataset. Batch correction was performed using a comprehensive integration algorithm^19^. Log normalization and scaling of features in the dataset was performed prior to principal component dimensionality reduction, clustering, and visualization using UMAP. Generally 15 PCAs were used in each analysis and resolution was set at 0.6. Differentially expressed genes and identification of cluster or cell type specific markers were identified using a Wilcoxon rank sum test between each defined group. P-value adjustment was performed using Bonferroni correction based on total number of genes in the aggregated dataset. The monocle library in R was used to determine pseudotime trajectories in separate cell subpopulations. Analysis of differential genes between clusters were performed using Ingenuity Pathway Analysis software.

### RNAseq analysis of human tumors

Collapsed RNA-Seq data from 123 human medulloblastoma tissues were obtained through data release V13 of the OpenPBTA project (github.com/AlexsLemonade/OpenPBTA-analysis), a global open science collaborative efforts of the Children’s Brain Tumor Tissue Consortium, Pediatric Neuro-oncology Consortium, Alex’s Lemonade Stand Foundation’s Childhood Cancer Data Lab, and the Center for Data-Driven Discovery in Biomedicine at the Children’s Hospital of Philadelphia. Microenvironment Cell Populations-counter (MCP-Counter) method from the R package immunedeconv^16^ was used to deconvolute the tumor microenvironment of 123 human medulloblastoma RNA-Seq samples consisting of four molecular subtypes i.e. Group3 (n = 15), WNT (n = 11), Group4 (n = 67) and SHH (n = 31). MCP-Counter represents cell type enrichment as abundance scores that are correlated to actual cell type proportions. To visualize the subtype specific enrichment, we created a heatmap of average immune scores per cell type across each molecular subtype (Code availability: https://github.com/d3b-center/Dang_MB_2020).

BRETIGEA^17^ method (https://github.com/andymckenzie/BRETIGEA) was used to find surrogate proportion variables (SPV) of brain cells astrocytes (ast), endothelial cells (end), microglia (mic), neurons (neu), oligodendrocytes (oli), and oligodendrocyte precursor cells (opc), derived from each of human, mice, and combination human/mouse data sets. In addition to that, we added monocyte marker genes from prior published work^14^ and monocyte marker genes from xCell^20^ to calculatesurrogate proportion variables for monocyte cell types in brain samples. The results should be considered as preliminary data which needs further validation by correlating SPV to true monocyte cell proportions in control datasets. We ran function findCells() using SVD method to calculate SPVs and all 1000 marker genes for brain cell types provided in BRETIGEA package along with 317 genes for monocyte. All cell type SPVs were then plotted for each sample as stacked bar plots.

### Immunohistochemistry

Whole mouse brain tissues were fixed in 4% formaldehyde for 7 days and standard paraffinization was performed. Sections were cut to 5 um thickness. Sections were rehydrated in xylene and serial ethanol concentrations. Antigen retrieval was achieved with sodium citrate buffer (ph7 or 9) in a pressure cooker. Sections were incubated overnight at 4°C with primary antibody. Anti-mouse primary antibodies used include the following: IbaI (Wako Chemicals, 019-19741), F4/80 (Life Technologies, MF48000), CD8 (Abcam, ab203035), and Ly6G (StemCell Technologies 60031). Tissues were then incubated with secondary anti-rabbit biotinylated (Vector Labs, BA-1000) or anti-rat biotinylated secondary antibody (Vector Labs, BA-4001) for 30 minutes at room temperature. Signal was amplified with avidin/biotin ABC complex (Vector Labs, PK-6102) and stained with DAB substrate chromogen (DAKO, 2016–10). NIS Elements BR 3.0 software was used to capture and analyze the images. Quantification of positive CD8, F4/80 or Ly6G cells were done in Imagescope. Counts are averages of 3 animal per treatment group. 3 sections spaced at least 100 microns apart were averaged for each animal.

### RNA in situ hybridization

For Chromogenic InSitu Hybridization (CISH) staining fresh slides were section, air dried, and baked within 48hrs of staining. Staining was performed on a Bond RXm automated staining system (Leica Biosystems). For dual CISH probe staining <u>Mm-Siglec</u> and <u>Mm-Clec12a</u> probes (Advanced Cell Diagnostics, <u>528248/514358-C2</u>) were used along with the RNAscope 2.5 LS Duplex Reagent Kit (Advanced Cell Diagnostics, 32240). Standardized protocols from Advanced Cell Diagnostics were used without modifications. For dual CISH + IHC Mm-Siglec or Mm-Clec12a probes (Advanced Cell Diagnostics 528248/514358) were used along with Iba 1 antibody (Wako 019-19741) at a 1:1K dilution with no additional retrieval steps. For the IHC portion a Bond Refine staining kit (Leica Biosystems, DS9800) was used with a standard protocol minus the peroxide blocking step which was deleted. After staining slides were air dried, coverslipped, and scanned at 40x magnification with an Aperio CS-O slide scanner (Leica Biosystems).

### Flow cytometry

Whole cerebellar tissue were minced and cells were dissociated with collagenase B and DNase I for 45 minutes at 37C and filtered through 70uM cell strainer. Samples were incubated for 10 minutes with CD16/32 Fc Block (BD Biosciences, 553142) and stained on ice with primary-fluorophore conjugated antibodies for identification of cell populations by FACS. Flow cytometry was peformed on an LSR II Flow Cytometer (BD Biosciences) and analyzed using FlowJo software (Treestar). Antibodies used include the following: CD45 (BioLegend, clone 30-F11), CD11b (BioLegend, clone CBRM1/5), Ly6C (BioLegend, clone HK1.4), Ly6G (BioLegend, 1A8), CD8 (BioLegend, 53.6.7), CD3 (BioLegend, 17A2).

### T cell suppression assay

Mouse splenic T cells were isolated from Ptch1^*+/*-^:Tp53^-/-^ mice using Pan T cell isolation Kit (Miltenyi Biotec). 4 × 10^4^ mouse T cells were labeled with CFSE (Life Technologies, C34554) and cultured for 3 days at 37C with 1ul of αCD3/28 bead (Thermo Fisher Scientific, 11131D) and 15U recombinant human IL-2 (Peprotech, Inc. 200-02). CD45+, Ly6Chi cells isolated from tumors post-radiation were sorted on MoFlo Astrios at the Children’s Hospital of Philadelphia Flow Cytometry Core Laboratories. T cell proliferation was determined by measuring CFSE signal in CD45^+^CD3^+^CD8^+^ cells.

### Statistical analysis

Statistical analyses of data were carried out using the unpaired two-tailed Student’s t-test for comparison between two experimental groups. Wilcoxon signed-rank test was used in the BRETIGEA human data analysis.

