## Supplementary figures for "Single-cell transcriptomic profile reveals macrophage heterogeneity in medulloblastoma and their treatment-dependent recruitment"

### Supplement Figure 1

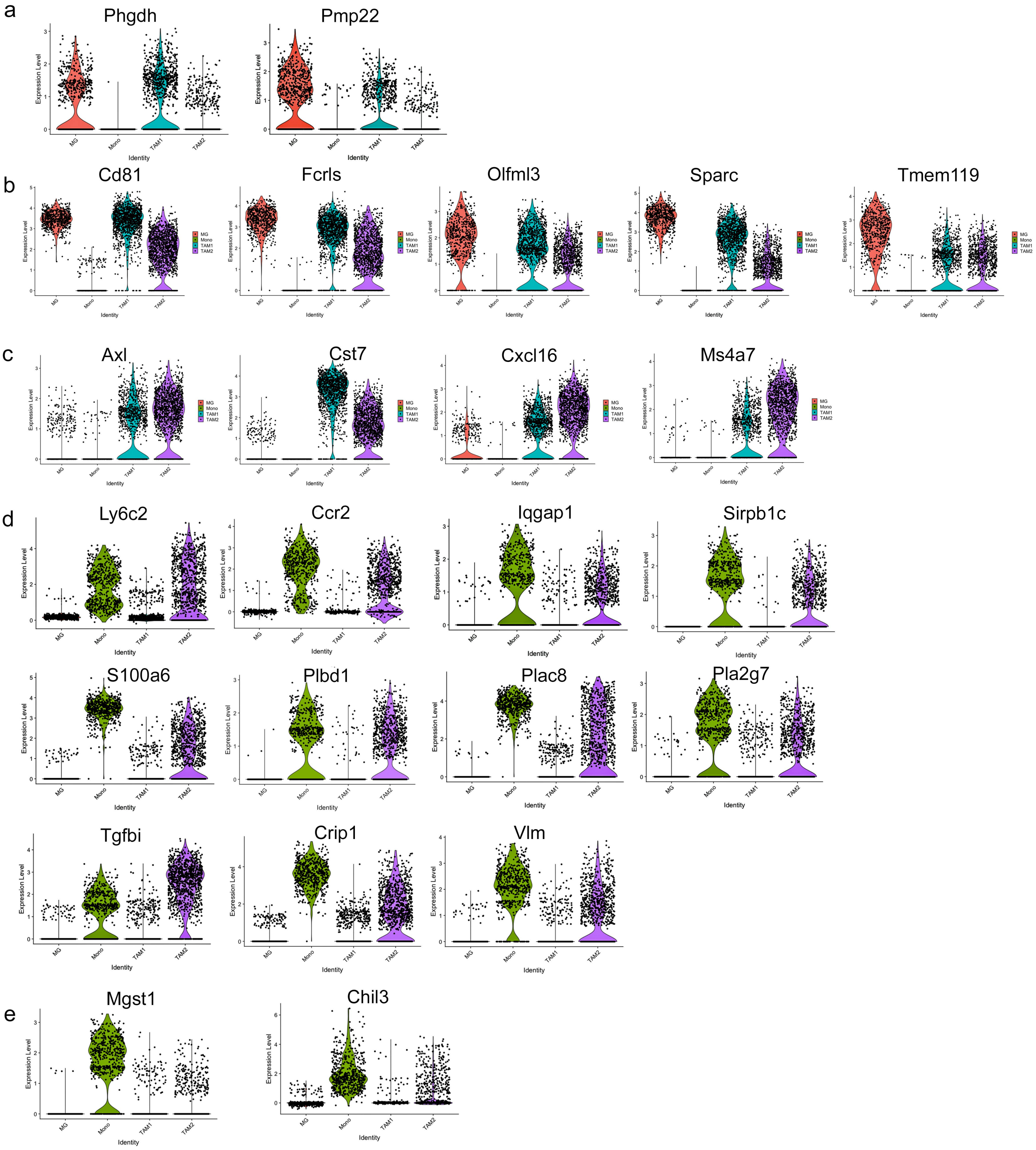

#### Supplement Figure 2

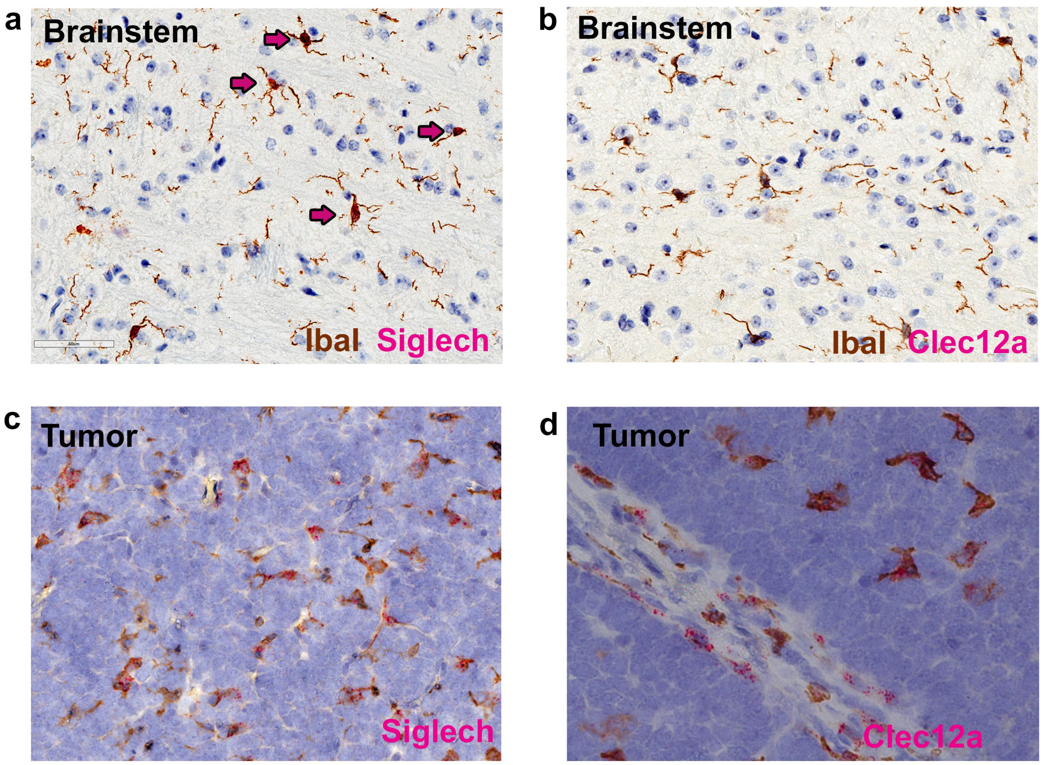

Supplementary Figure 3

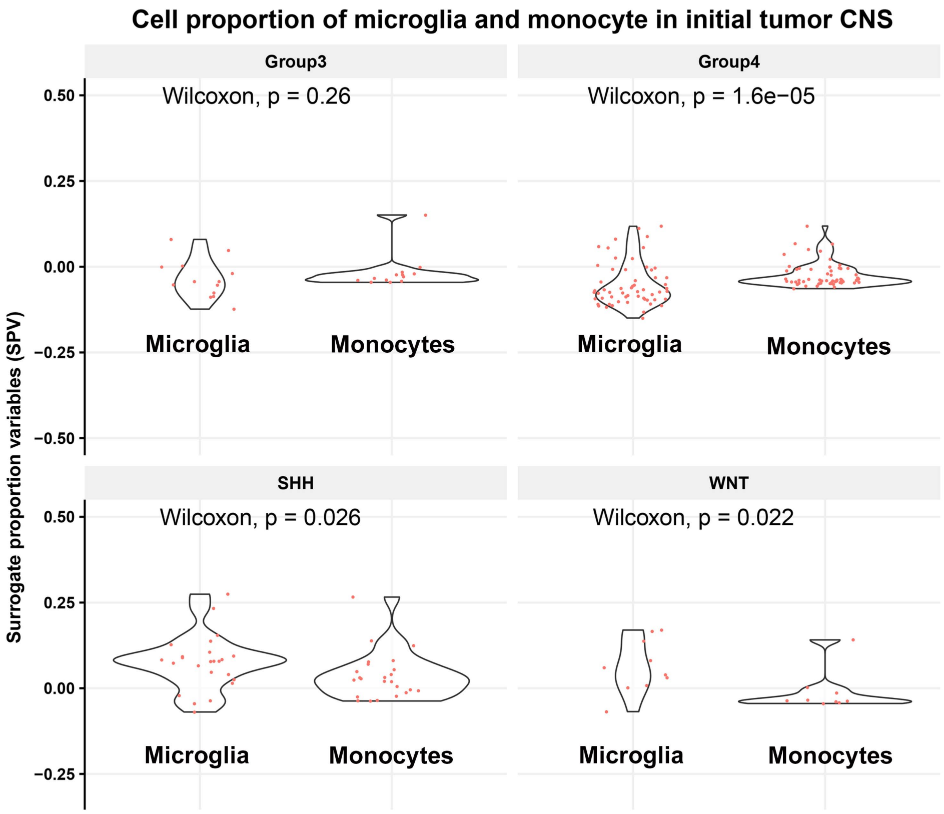

#### Supplement Figure 4

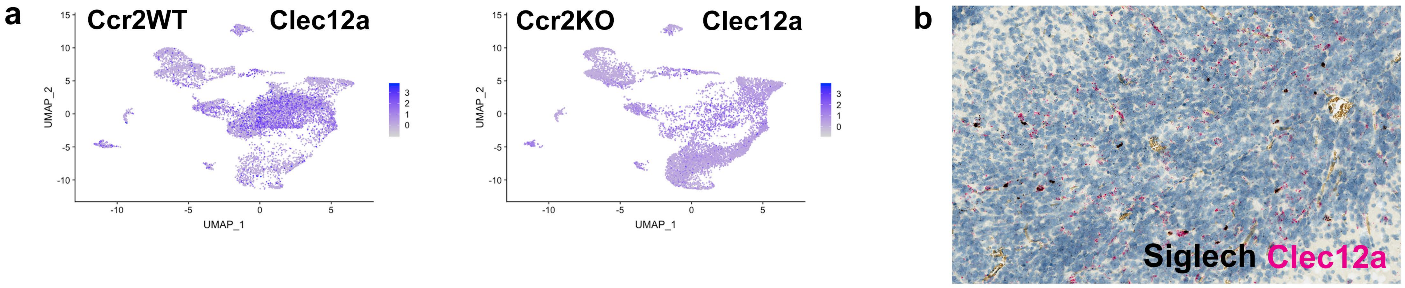

Supplementary Figure 5

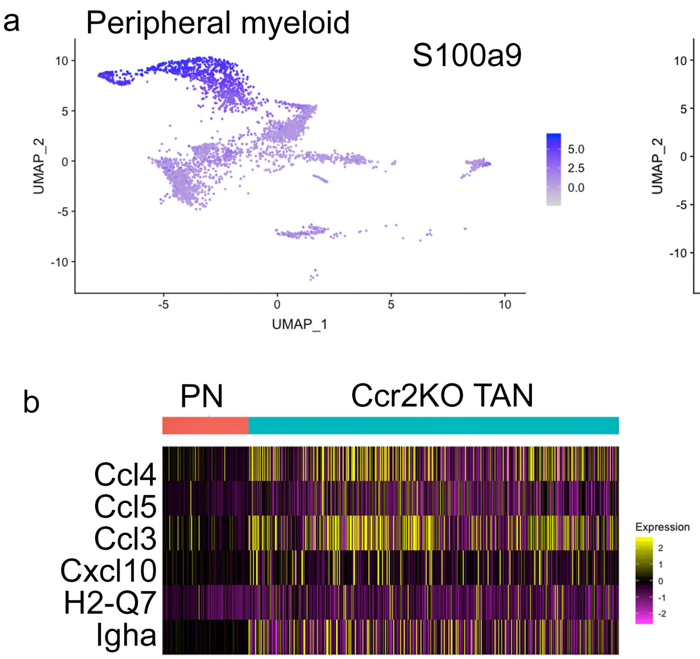
